# Fast and Robust Deconvolution of Tumor Infiltrating Lymphocyte from Expression Profiles using Least Trimmed Squares

**DOI:** 10.1101/358366

**Authors:** Yuning Hao, Ming Yan, Yu L. Lei, Yuying Xie

## Abstract

Gene-expression deconvolution is used to quantify different types of cells in a mixed population. It provides a highly promising solution to rapidly characterize the tumor-infiltrating immune landscape and identify cold cancers. However, a major challenge is that gene-expression data are frequently contaminated by many outliers that decrease the estimation accuracy. Thus, it is imperative to develop a robust deconvolution method that automatically decontaminates data by reliably detecting and removing outliers. We developed a new machine learning tool, <u>F</u>ast <u>A</u>nd <u>R</u>obust <u>DE</u>convolution of <u>E</u>xpression <u>P</u>rofiles (FARDEEP), to enumerate immune cell subsets from whole tumor tissue samples. To reduce noise in the tumor gene expression datasets, FARDEEP utilizes an adaptive least trimmed square to automatically detect and remove outliers before estimating the cell compositions. We show that FARDEEP is less susceptible to outliers and returns a better estimation of coefficients than the existing methods with both numerical simulations and real datasets. FARDEEP provides the absolute quantitation of each immune cell subset in addition to relative percentages. Hence, FARDEEP represents a novel robust algorithm to complement the existing toolkit for the characterization of tissue-infiltrating immune cell landscape. The source code for FARDEEP as implemented in R is available for download at https://goo.gl/SqGKuo.

## 1. Introduction

The immune system constitutes a primary host defense mechanism against malignancy. Somatic mutations can be detected by surveying antigen-presenting cells (APC), which facilitate the expansion of tumor-specific effector T cells. But these effectors rapidly become functionally exhausted upon entry into the immunosuppressive tumor microenvironment (TME), with high expression levels of the inhibitory immune checkpoint receptors (ICR). Blockade of ICR has generated broad enthusiasm in the clinics thanks to its potential in reinvigorating the pre-existing T-cell pool, but the majority of cancers are hypoimmunogenic with coordinated mechanisms to exclude APC and effectors, which drives resistance to ICR blockade immunotherapy. Thus, combinatorial strategies are essential to prime the immune system and sensitize cold cancers to ICR blockade. The success of these emerging strategies depends on the precise identification of cold cancers and selection of personalized treatment protocols.

Immunogenomics presents an unprecedented opportunity to stratify patients based on cancer immune infiltrates and complement the current TNM staging system. Multiple clinical studies highlight the prognostic significance of certain immune cell subsets in the TME (Balermpas *et al.*, 2014, 2016; Nguyen *et al.*, 2016; Pages *et al.*, 2009; Wolf *et al.*, 2015; Lei *et al.*, 2016). Immunohistochemistry (IHC)-based immunoscores, which quantify the number of CD8^+^ cytotoxic T lymphocytes and CD45RO^+^ memory T cells, show better prognostic potential than conventional pathological methods in colon cancer patients (Galon *et al.*, 2006; Mlecnik *et al.*, 2016). Hence, harnessing the composition of intra-tumoral immune cell infiltration is a highly promising approach to identify cold tumors. The current IHC immunoscoring approach has two limitations. First, the interpretation of immune cell subsets varies among pathologists and institutions, thus lacking a consistent standard for the scoring practice. Second, only a limited number of biomarkers can be assessed simultaneously, which prevents a comprehensive annotation of the immune contexture in the TME. Hence, robust methods for genome data-informed cell type quantitation are in urgent need. A significant technical obstacle is that the efficacy and accuracy of gene expression deconvolution are limited by the large number of outliers, which are frequently observed in tumor gene expression datasets (Jiang *et al.*, 2004). The first step towards enhancing the overall gene deconvolution algorithms is to improve methods for outliers identification and processing. In this study, we report a novel FAst and Robust DEconvolution of Expression Profiles (FARDEEP) method that significantly improves the estimation of coefficients.

Let *y_i_* be the observed expression value for the *i*th gene; *x_i_*, a *p*-dimensional vector, be the expected expression of the *i*th gene for the *p* different cell types. The gene-expression deconvolution problem can be formulated as equation (0.1),

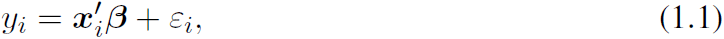

where *β* ∈ ℝ^p^ is an unknown parameter corresponding to the compositions of *p* types of cell, and *∊_i_* is a noise term with a mean of 0. Several methods were proposed to solve this deconvolution problem. To enforce the non-negativity of *β*, several groups have proposed using the Non-Negative Least Square (NNLS) and Non-negative Maximum Likelihood (NNML) frameworks to solve (1.1) (Lawson and Hanson, 1995; Gong *et al.*, 2011; Gong and Szustakowski, 2013; Li *et al.*, 2016; Mackey *et al.*, 1996; Qiao *et al.*, 2012). Additionally, the gene expression of each cell may vary depending on its microenvironment and other factors, which will lead to a biased estimation. To address this issue, Microarray Microdissection with Analysis of Differences (MMAD) incorporates the concept of the effective RNA fraction, and estimates coefficients using a maximum likelihood approach (Liebner *et al.*, 2014). To further adapt deconvolution to high-dimensional settings, Altboum *et al.* (2014) proposed a penalized regression framework, Digital Cell Quantifier (DCQ), to encourage sparsity for the estimated *β* using elastic net Zou and Hastie (2005). CIBERSORT introduces *ν*-support vector regression (*ν*-SVR) to enhance the robustness of gene expression deconvolution. The method performs a regression by finding a hyperplane that fits as many data points as possible within a tube whose vertical length is a constant *ε* (Newman *et al.*, 2015). The *ε*-tube provides a region in which estimation errors are ignored. However, CIBERSORT does not include an intercept in the model to capture contributions of other cell types. Additionally, to increase the computational efficiency CIBERSORT apply z-normalization to the data before fitting the regression, which introduces estimation bias. Discussion of the effect of Z-score normalization for CIBERSORT is included in Supplementary material. The number of effector immune cells is a critical factor underpinning the efficacy of immune killing. Hence, a robust method that determines both the distribution and absolute volume of tumor-infiltrating lymphocytes (TILs) will substantially improve the current gene deconvolution pipeline.

To handle contaminated gene expression data and provide absolute cell abundance estimation, we developed a robust method based on the Least Trimmed Square (LTS) framework (Rousseeuw, 1984; Rousseeuw and Leroy, 1987). LTS finds *h* observations with smallest residuals, and the estimator 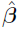 is the least squares fit over these *h* obeservations. LTS is a NP hard problem, and Rousseeuw and Driessen (2006) proposed a stochastic FAST-LTS algorithm. Nevertheless, it may give a suboptimal fitting result and get much slower when the sample size and dimension of variables become larger and higher since its accuracy relies on the initial random *h*-subsets and the number of intial subsets. When *n* is the sample size and *p* is the number of coefficients, *h* is suggested to be the smallest integer that is not less than (*n* + *p* + 1)*/*2 to remove as many outliers as possible while keeping a unbias result. Using the information of only half of the data reduces the power of the estimator because the amount of outliers in real case cannot be presumed and can be small. Xu *et al.* (2017) proposed an adaptive least trimmed square which is not limited to the randomness of initial subset but only applied in binary dataset. In this study, we extend the adaptive least trimmed square to introduce a model-free and efficient method, which could find the number of outliers automatically based on LTS. As evidence of high fidelity and robustness, the performance of FARDEEP exceeds that of the existing tools in simulated and real world datasets.

## 2. MATERIALS AND METHODS

### 2.1. Model formulation

The usual linear deconvolution model can be expressed as below,

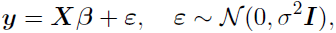

where *y ∈* ℝ^q^ is the observed expression data for *q* immune subset signature genes, *X ∈ ℝ^q×p^* denotes a mean gene expression signature matrix for *p* different cell types, *β ∈ ℝ^p^* contains each unknown cell type abundance, and *ε ∈* ℝ*^q^* is a vector of random errors. To incorporate outliers, we propose the following model

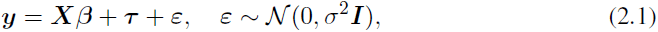

where parameter 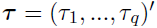 is a sparse vector in ℝ^q^ with *τ_i_* ≠ 0 indicating the *i*th gene is an outlier.

Under the formulation of (2.1), let 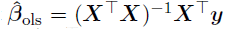 be the Ordinary Least Square (OLS) estimate and ***H*** = ***X***(***X**^T^**X***)*^−1^ **X**^T^* be the projection matrix. The residuals *r* = (*r*_1_*, …, r_q_*) using OLS could be formulated as

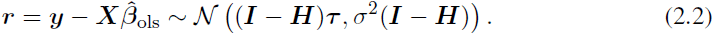

### 2.2. Adaptive least trimmed square

From (2.2), the residuals, *r_i_* with the corresponding *τ_i_* ≠ 0, would deviate from zero, which suggests that the set of outliers can be identified through thresholding as follows

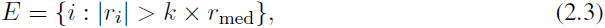

where *E* is the set of detected outliers, *k* is a tuning parameter controlling the sensitivity of the model, and *r*_med_ is the median of 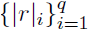. We denote the number of elements in set *E* as *|E|* and let *N* be the number of true outliers in the data. First, we can use least squares and formula (2.3) to obtain a rough estimate of *E* denoted as *Ê*. Let the cardinality of *Ê* be 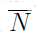. Since the model at this moment is inaccurate with contamination of outliers, 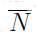 is an overestimation of *N* which can be used to get an underestimate via *<u>N</u>* = 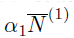 with α_1_ *∈* (0, 1). With *<u>N</u>*, we can then update the least square fitting after removing the *<u>N</u>* samples with the largest absolute value of residuals and obtain an improved estimate of *E* and the corresponding 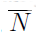. We can improve the model by repeating the procedure, but we need to increase the underestimate of outliers, *<u>N</u>*, by a factor of α_2_ with α_2_ *>* 1 for each iteration to force the convergence between 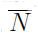 and *<u>N</u>*. In sum, for the *j*th iteration, we update 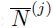 by

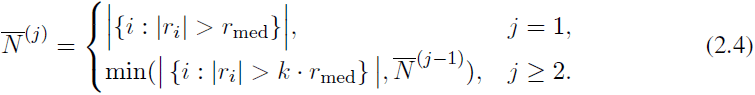

where the min(*·, ·*) operator guarantees that 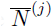, an overestimation of *N*, is non-increasing after each iteration. Similarly, we update *<u>N</u>*^(*j*)^ through

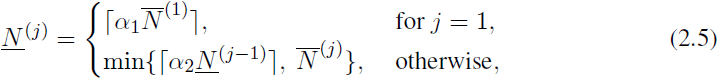

where *⌈x⌉* means the ceiling of *x ∈* ℝ, α_1_ *∈* (0, 1) is used to obtain a lower bound for *N* in the first step, α_2_ *>* 1 guarantees the monotonicity of *<u>N</u>*^(*j*)^, and the min(*·, ·*) operator makes sure *<u>N</u>* ^(*j*)^ is smaller than 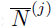. Then we update 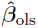 after removing *<u>N</u>*^(*j*)^ outliers, and the algorithm stops when *<u>N</u>* and 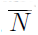 converge.

Hence, we hereby report a novel approach, coined as adaptive Least Trimmed Square (aLTS), to automatically detect and remove contaminating outliers. Our aLTS is an extension of the iterative LTS algorithm proposed by Xu *et al.* (2017) which is designed for binary output such as comparison between two images or videos.

### 2.3. FARDEEP

Because the amount of cell types is always non-negative, we replaced the OLS regression in the procedure of aLTS with non-negative least square regression. By applying aLTS to the deconvolution model (0.2) and solving the following problem,

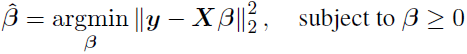

using Lawson-Hanson algorithm (Lawson and Hanson, 1995), we developed a robust tool, FARDEEP, for cellular deconvolution (Algorithm 1).

#### Algorithm 1 FAst and Robust DEconvolution of Expression Profiles

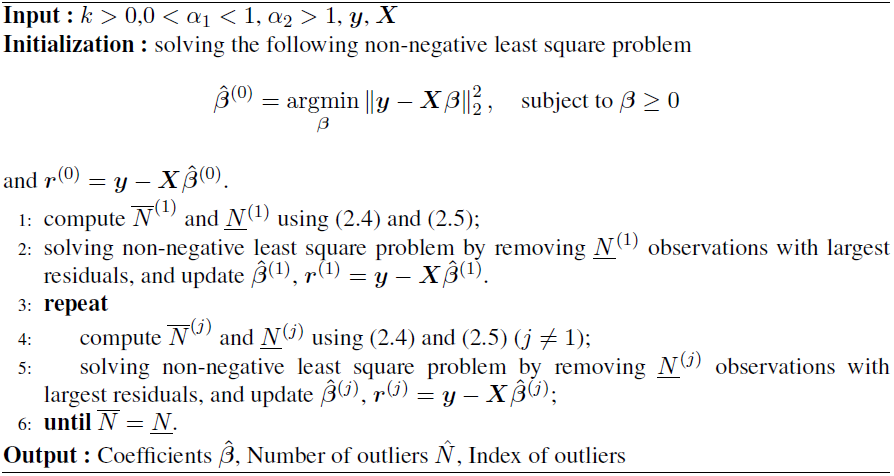

One unique advantage of FARDEEP is that it is fast and guarantees to converge within finite steps, which is summarized in the following theorem.

#### Theorem 1. Algorithm 1 (FARDEEP) stops in no more than j^*^ steps, where

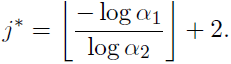

Here ⌊·⌋ is the largest integer that is less than or equal to x.

*Proof.* It follows from the fact that the sequence 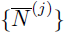> is non-increasing, and *{<u>N</u>*^(*j*)}^ is a geometrically increasing sequence that is bounded by the smallest component of 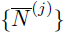. Specifically, assume that *j^*^* steps have been taken in FARDEEP, then *j* has approached *j^*^ −*1, and *<u>N</u>*^(*j*)^≥ α_2_*N*^(*j−*1)^ for 0 ≤ *j* ≤ *j^*^* − 1, so

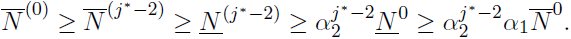

which leads to the result.

### 2.4. Parameter tuning

There are three tuning parameters *k*, α_1_, and α_2_ in FARDEEP. Since α_1_ is only used in the first iteration, a relatively small α_1_ is preferred to ensure that FARDEEP does not remove too many outliers at the first step. In practice, FARDEEP is not sensitive to different values of α_1_, and α_2_. However, *k* controls the number of outliers in each iteration and is critical for the performance of FARDEEP. *k* is fine tuned on a case-by-case basis to preserve meaningful fluctuations of gene expression levels. Effects for different tuning parameters are shown in Supplementary Table S1. Cross-validation is not advised since the test group may contain outliers that influence the accuracy of the tuning result. Instead, we applied the Bayesian Information Criterion (BIC) and assume that the errors follow a log-normal distribution instead of a normal distribution among gene expression datasets as suggest by Beal (2017). We define the modified BIC referring to the setting of She and Owen (2011):

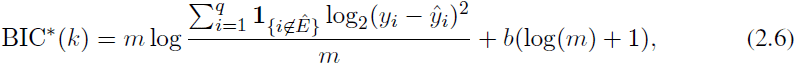

where *Ê* being the set of detected outliers, *b* is number of parameters and equals 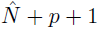 with 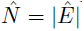 being the number of outliers, and *m* equals *q −*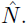. Then, we choose the value of *k* associated with the smallest BIC^*^.

## 3. RESULTS

To test the performance of FARDEEP, we compared our approach with the existing methods using numerical simulations and real datasets. We first used LM22 as the signature matrix that was generated by Newman *et al.* (2015). This signature matrix contains 547 genes that could be used to accurately deconvolve 22 human hematopoietic cells. We use the sum of squared error (SSE) and coefficient of determination denoted as R-squared (*R*^2^) as the metrics:

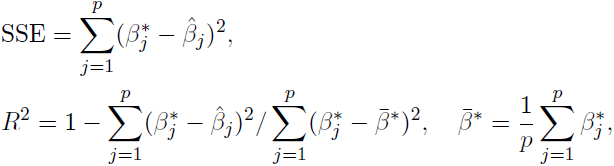

where *β^*^* is the true amount of cells, and 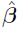 is the estimate. Here, we use the coefficient of determination instead of the Pearson correlation because we are focusing on the estimation accuracy for the absolute not the relative amount while the Pearson correlation can only measure the strength of linear relationship. For instance, if the true amount is β^*^ = (0.∈, 0.4, 0.6)*^′^* and the estimate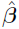 is = (0.1, 0.0.2, 0.3) which is not a good estimate, the Pearson correlation is exactly 1 treating it as a perfect estimate. On the other hand, the corresponding coefficient of determination is *−*0.75 suggesting the poor performance of the results.

### 3.1. In silico simulation based on leukocyte gene signature matrix file

We randomly generated the amount of different cells from interval [0, 1]. Notably, the sum of cell abundance is not necessarily 1. The measurement errors were sampled from 2*^N^*(0, (0.1 log_2_(*s*))^2^). To incorporate outliers, we randomly selected *i/*50 of the data and replaced them with data drawn from 2*^N^*(10, (0.3 log_2_(*s*))^2^) where *i* = 1, 2*, …*, 25 and *s* is the standard deviation of original mixtures (Newman *et al.*, 2015).

We repeated the procedure for nine times and compared the coefficients estimator using the SSE and coefficient of determination (*R*^2^). By comparing the results of FARDEEP, CIBER-SORT (without converting to percentage), NNLS, PERT and DCQ, we found that the SSE range for FARDEEP is 3.53 *×* 10*^−2^* to 2.92 *×* 10*^−4^* and *R*^2^ keeps being 1 regardless of the number of outliers. Other methods show significantly larger SSE and smaller *R*^2^ (Supplementary Table S2).

### 3.2. In silico simulation with heavy-tailed error

To test the robustness of FARDEEP under heavy-tailed random error, we simulated a new dataset following the setting in She and Owen (2011); Alfons *et al.* (2013). The observations were generated based on the linear regression model (2.1). The predictor matrix is X = (*x*_1_*, …, x_n_*)*^′^* = UΣ ^½^, where *U_ij_* ˜ *ρ U*(0, 20) and Σ*_ij_* = *ρ^I^{*i*≠*j*}* with *ρ* = 0.5. Consider the proportion of outliers *f ∊ {*5%, 10%, 20%, 30%*}*, sample size *n* = 500, and number of predictors *p* = 20, we randomly added outliers to the simulated data as follows:

i) Vertical outliers: we generated a zero vector *τ*, and randomly selected *nf* elements in *τ* to be the outlier terms. Then, we replaced these elements by *nf* samples which were generated from non-central t-distribution with 1 degree of freedom and a non-centrality parameter of 30.

ii) Leverage points: we took 20% of the contaminated data in i) as leverage points, that is, replacing the corresponding predictors by the samples from *N* (2max(*X*), 1). The coefficients *β_j_* and random errors ∊_*i*_ were respectively sampled from *U*(0, 1) and t-distribution with 3 degrees of freedom, where *j* = 1*, …, p* and *i* = 1*, …, n*. Based on the framework above, the dependent variable could be obtained by

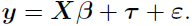

We simulated each model 50 times. As shown in Figures 1 and 2, FARDEEP outperforms other methods, evidenced by the SSE and *R*^2^ values.

**Figure 1.**
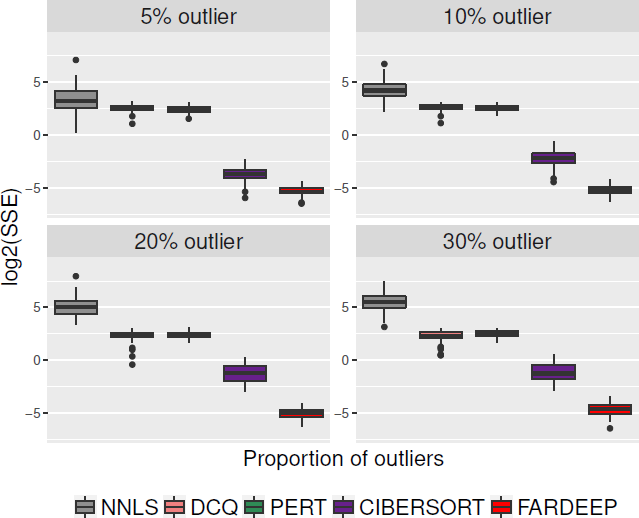
SSE of coefficients for different approaches. We simulated different percentage of outliers (*{*5%, 10%, 20%, 30%*}*) and compared the SSE for coefficients applying NNLS, DCQ, PERT, CIBERSORT and FARDEEP.

**Figure 2.**
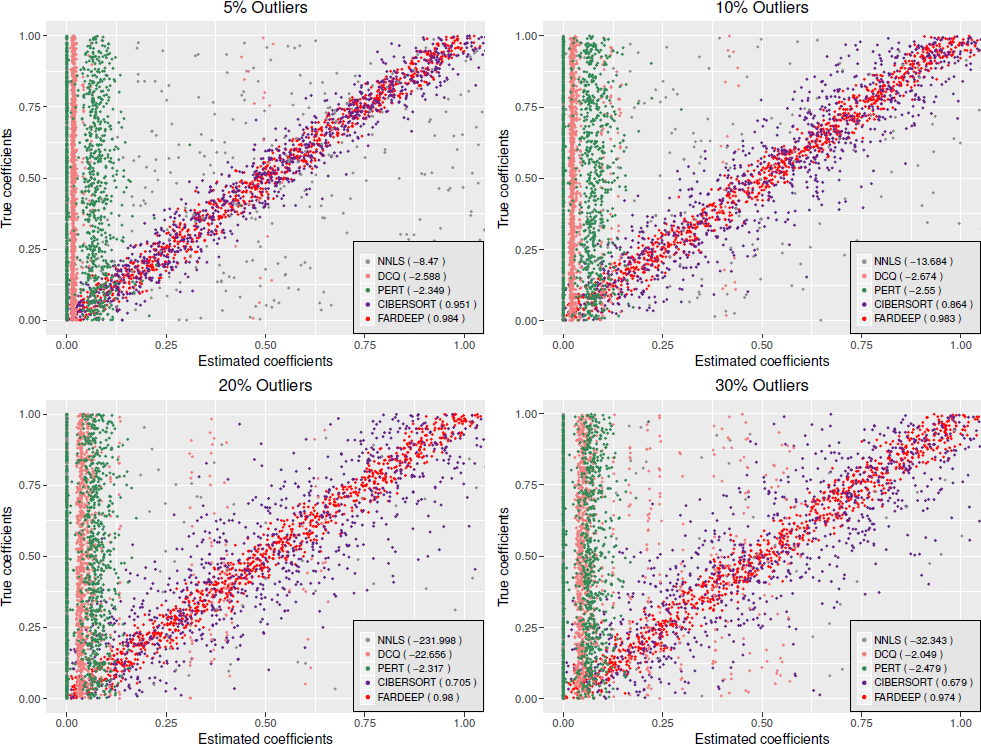
Compare the accuracy of different deconvolution approaches (the values in parentheses are *R*^2^). Based on {5%, 10%, 20%, 30%*}* percentage of outliers, we computed *R*^2^ to evaluate how well are the estimators fit for a straight line 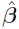 = β. PERT and DCQ fail to estimate the coefficients accurately (*R*^2^ are negative since the SSE are too large), FARDEEP shows the best performance in each case.

To check FARDEEP’s accuracy of outlier detection, we simulated {5%, 10%, 20%, 30%, 40%*}* outliers using the same method as above for a model with both normal distributed noise and heavy-tailed noise. Tuning parameter *k* using the modified BIC in (2.6), we show that *k* decreases when the amount of outliers becomes larger. The true positive rates always stay around 1, indicating that the tunning of *k* is highly effective (Table 1).

**Table 1.**
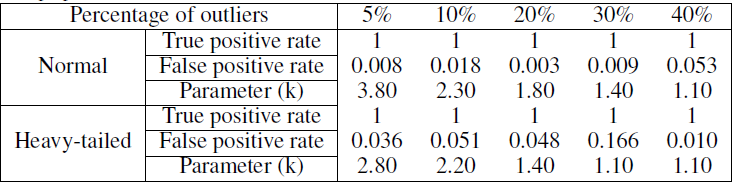
Tuned *k* for FARDEEP with the adjusted BIC. We simulated normal distributed error and heavy-tailed error respectively for different proportion of outliers and computed true positive rate and false positive rate to evaluate the tunning result.

### 3.3. Synthetic dataset

We used the cell line dataset GSE11103 generated by Abbas *et al.* (2009) that contains gene expression profiles of four immune cell lines (Jurkat, IM-9, Raji, and THP-1) and four mixtures (MixA, MixB, MixC, and MixD) with various ratios of cells. Before analysis, we quantile normalized the mixture data for 54675 genes and downloaded the immune gene signature matrix with 584 genes from CIBERSORT website. Then, we applied five deconvolution methods, including FARDEEP, CIBERSORT (without converting to percentage), DCQ, NNLS and PERT, to calculate the sum of squared errors of the estimated amount of the four immune cell lines. We also compared with CIBERSORT absolute mode, which is a beta version in CIBER-SORT website (Supplementary Figure S1). Since the CIBERSORT absolute mode is a beta version and leads to suboptimal results compared with CIBERSORT, we only focused on the comparisons with CIBERSORT. We show that FARDEEP gives the smallest SSE and the largest *R*^2^, which indicates the most accurate result (Figure 3).

**Figure 3.**
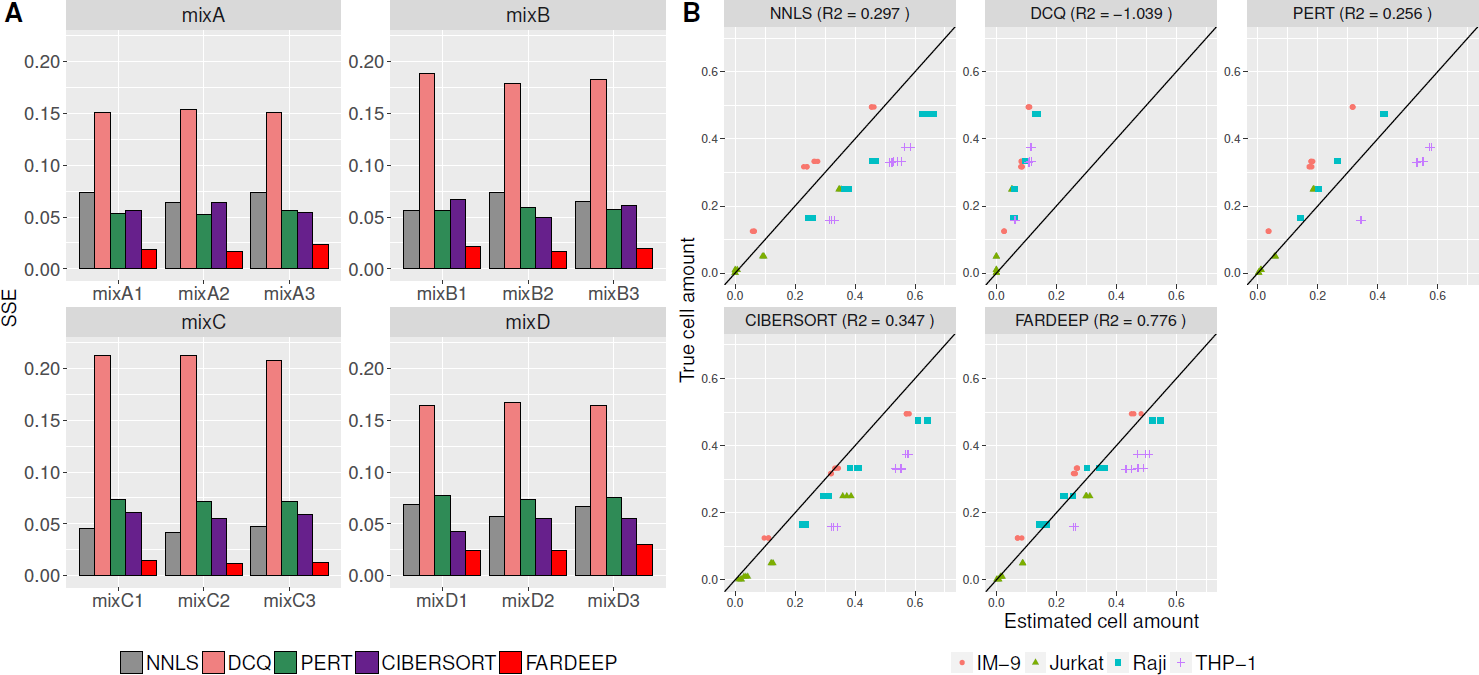
Applying different deconvolution approaches on the gene expression data of IM-9, Jurkat, Raji, THP-1 and the mixture of these four immune cell lines with known proportion (MixA, MixB, MixC, MixD). All of the mixtures were performed and measured in triplicate. (A) SSE of coefficients for FARDEEP, CIBERSORT, NNLS, PERT, DCQ. (B) Amount of cell lines estimated from different deconvolution approaches vs. amount of cell lines truly mixed, and using R-squared to evaluate the performance of each method.

### 3.4. Synthetic dataset with added unknown content

Both CIBERSORT and FARDEEP are robust deconvolution methods and show advantages in the above datasets, we next sought to compare their performances on mixtures with unknown content. We followed the simulation setting proposed by Newman *et al.* (2015) and downloaded the mixture dataset and the signature gene file from CIBERSORT website. The mixture file was constructed from the four immune cell lines data, as mentioned in the previous section, and a colon cancer cell line ( average of GSM269529 and GSM269530 in GSE10650). Cancer cells were mingled into immune cells at different ratios{0%, 30%, 60%, 90%*}*. Noise was added by sampling from the distribution 2*^N^* ^(0,(*f* log_2_(*s*))^2^)^, in which *f ∊{*0%, 30%, 60%, 90%*}* and *s* is the standard deviation of original mixtures. By applying FARDEEP and CIBERSORT (without converting to percentage) on 64 mixtures, we found that FARDEEP remains an accurate estimation, while the tumor contents skews the results of CIBERSORT with larger deviation from the the ground truth (Figure 4).

**Figure 4.**
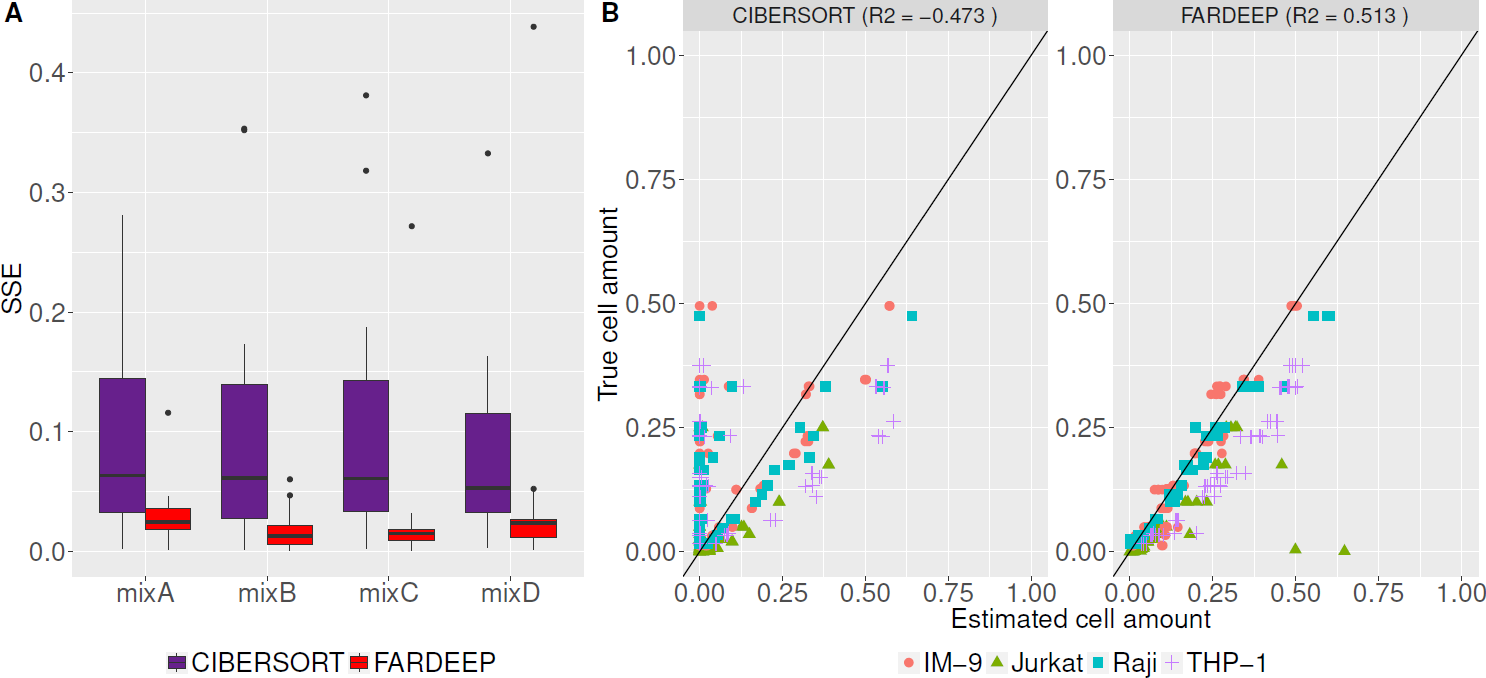
Comparing the performance of CIBERSORT and FARDEEP on mixtures with unknown content by SSE and *R*^2^. (A) SSE for various noise and amount of tumor contents on 4 different mixtures. (B) Amount of cell lines estimated by CIBERSORT and FARDEEP vs. amount of cell lines truly mixed.

### 3.5. Deconvolution performance on immune-cell-rich datasets

To evaluate the performance of FARDEEP in immune-cell-rich settings that are less affected by outliers, we downloaded and analyzed two gene expression datasets (GSE65136) from lymph node biopsy and peripheral blood used in Newman *et al.* (2015). In the two datasets, gene expression data were generated from Illumina BeadChip which is the same platform used for the signature matrix LM22, and the TILs of interest have abundance consists on average greater than 5%. These studies contains both gene expression and cell flow cytometry data measuring multiple immune cell subsets and normal tissues:

(i) surgical lymph node biopsies of follicular lymphoma patients;
(ii) purified B and T cells from the tonsils of five healthy controls;

Flow cytometry data in these studies, which are treated as ground truth, are in relative scales, thus we normalized the estimated parameters of each method to sum of 1 before comparison.

As shown in Figure 5A for case (i), FARDEEP outperformed CIBERSORT in terms of *R*^2^ and SSE, which is consistent with our findings with simulated datasets. For case (ii), we estimated the immune cell composition for purified B and T cells with purity level exceeding 95% and 98%, respectively. For purified B cells, CIBERSORT tends to return a non-zero estimates for T cell and a large proportion of other cell types, while FARDEEP gave almost all zero estimates for T cell and on average reduced the estimation error by 61%. Similarly, for the purified T cell, although CIBERSORT had a better performance compared to purified B cell, FARDEEP still improved significantly improve the estimation accuracy by reducing on average 48% of the estimation error (Figure 5B).

**Figure 5.**
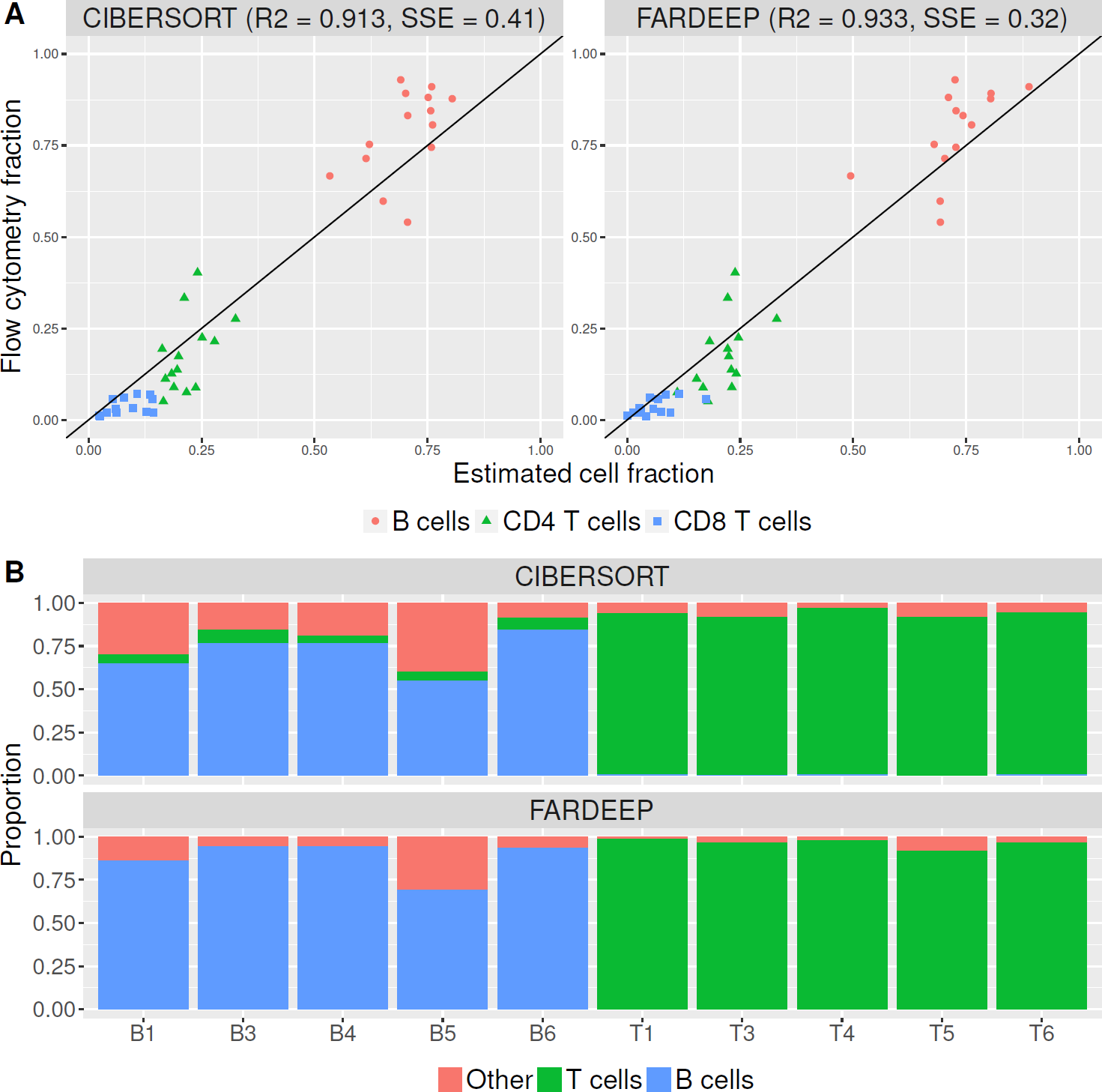
Performance assessment on real data with SSE and *R*^2^. (A) Follicular lymphoma dataset with estimation of B cell, CD8 T cell, and CD4 T cell using FARDEEP, and CIBERSORT. (B) Normal tonsil dataset with purified B cells and T cells.

Overall, FARDEEP and CIBERSORT are both robust methods, which generate generally comparable results in specimens that are rich in immune cells and contain minimal carcinoma cell contamination.

### 3.6. TCGA Ovarian Serous Cystadenocarcinoma dataset

However, the TME of solid carcinomas are far different from a lymph node biopsy or peripheral blood in that the highly variable gene expression in cancer cells challenges the accuracy of immune cell deconvolution. Hence, we next utilized survival and gene expression data (n = 514) of the ovarian cancer TCGA database to assess the prognostic relevance of different deconvolution methods. Using gene expression data (n = 514), we estimated the immunoscore using ESTIMATE proposed by Yoshihara *et al.* (2013) as well as TILs abundance using CIBERSORT and FARDEEP. Cold tumors typically harbor lower numbers of CD8^+^ T cells, *γ^δ^* T cells, M1-macrophages, and NK cells (Lei *et al.*, 2016; Qiao *et al.*, 2017; Binnewies *et al.*, 2018; Corrales *et al.*, 2016). Thus, we calculated the T_H_1/Tc1-skewed immune compartment by the summation of CD8^+^ T cells, *γ^δ^* T cells, M1-macrophages, and NK cells. Then, we partitioned the patients into two groups with equal size using the median of either the immunoscore (ESTIMATE) or T_H_1/Tc1-skewed immune compartment (CIBER-SORT and FARDEEP). We compared the survival curves between the two groups. As shown in Figure 6, FARDEEP most effectively separates patients into high-and-low risk groups with the smallest p-value (p-value = 0.0065). Recently, CIBERSORT website supports a beta-11 version of absolute mode for cell deconvolution, which report the absolute instead of relative abundance of TILs. We also included (CIBERSORT absolute mode) in this survival analysis and showed that it returned a slightly better results (p-value = 0.037) compared to the relative mode but still was dominated by FARDEEP (Figure S2). These results demonstrated that the absolute abundances of TILs could provide additional clinical-relevant information compared to the relative abundance. Additionally, these results suggest that FARDEEP can complement the current robust methods, such as ESTIMATE and CIBERSORT, for TIL deconvolution challenges in datasets.

**Figure 6.**
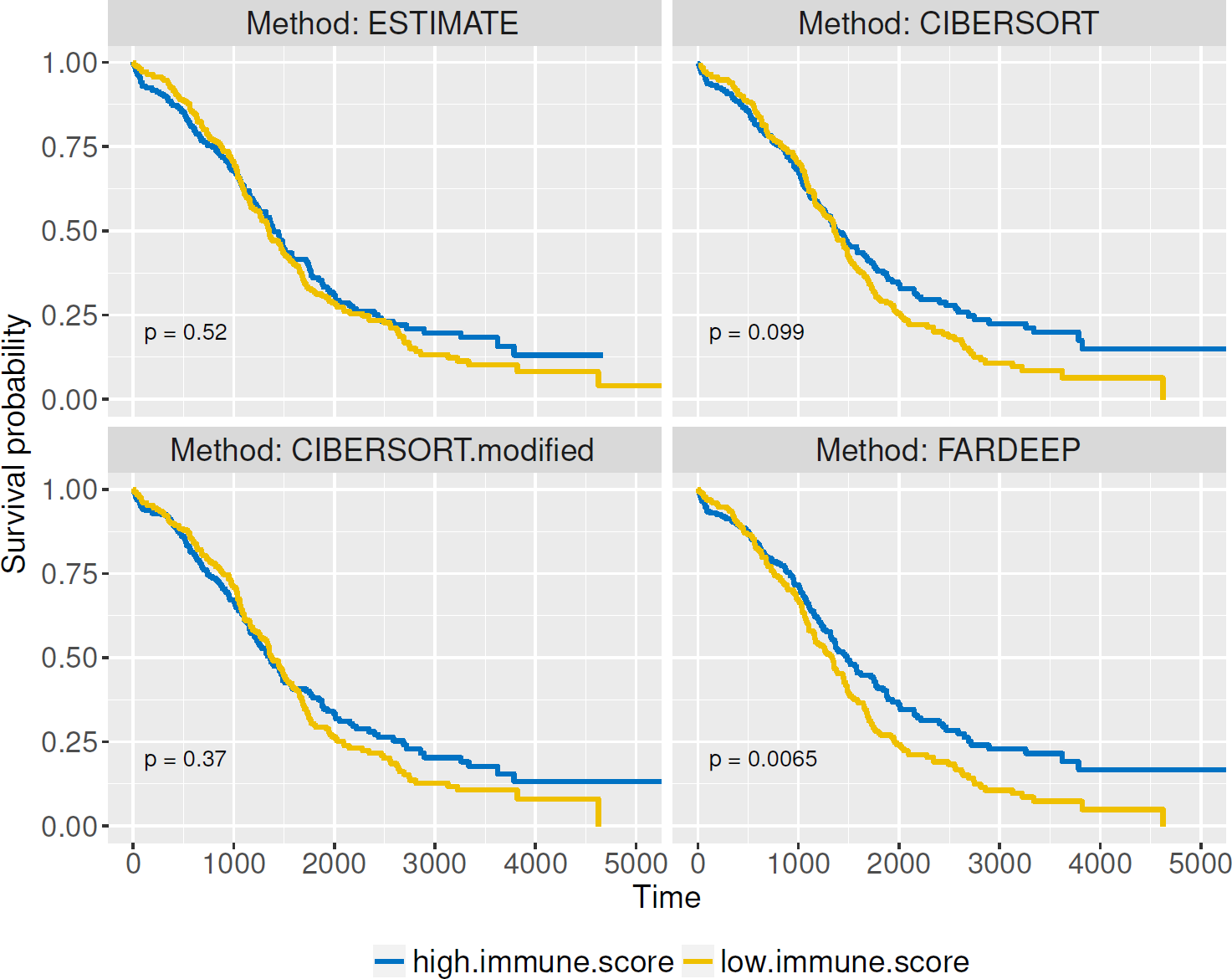
Kaplan-Meier survival curves are plotted based on ESTIMATE, FARDEEP- and CIBERSORT- assisted TIL profiling. Log-Rank test were applied to data for 514 patients with ovarian cancer classified into two groups according to whether immunoscore (ESTIMATE) or T_H_1/Tc1-skewed immune compartment (CIBERSORT and FARDEEP) was above or below the median.

## 4. DISCUSSION

Cancer immune microenvironment has emerged as a critical prognostic dimension that modulates patient responses to neoadjuvant therapy. However, the current clinical TNM staging system does not have a consistent method to stratify cancers based on their immunogenicity. Recent study shows that the RNA-Seq datasets of whole tumors contain valuable prognostic information to assess the cancer-immunity interactions (Newman *et al.*, 2015; Robinson *et al.*, 2017). But the current methods to extract immune signatures are susceptible to the frequent outliers in the datasets, leading to less effective identification of cold tumors. In this study, we developed a new machine learning tool, FARDEEP, to streamline the removal of outliers and increase the robustness of gene-expression profile deconvolution. Robustness is an indispensable feature to solve a problem of deconvolution because gene expression data are frequently contaminated by large amount of outliers. FARDEEP solves the deconvolution problem in a robust way because this tool evaluates all outliers across the datasets and then examines the true immune gene signature using non-negative regression. This feature is especially useful to analyze tumors with significant non-hematopoietic tumor components. Interestingly, although FARDEEP and the current robust methods can both tackle immune-cell-rich specimens such as lymph node and PBMCs, FARDEEP exhibits improved prognostic potential when dealing more complex datasets with significant carcinoma cell content.

Overall, here we show that FARDEEP is a powerful and rapid machine learning tool that outperforms existing robust methods for gene deconvolution in datasets with significant heavy-tailed noise. FARDEEP provides a new technology to interrogate cancer immunogenomics and more accurately map the immune landscape of solid tumors.

## 5. ACKNOWLEDGEMENTS

This work is supported by NIH grants R03 DE027399 (YLL and YX), R01 DE026728 (YLL) and R00 DE024173 (YLL), NSF grant DMS-1621798 (MY), the Michigan State University STEM Gateway Fellowship (YX), State Key Laboratory of Oral Diseases (SKLOD) Open Fund of China (YX), University of Michigan Rogel Cancer Center Research Grant (YLL) and POM Clinical Research Supplement (YLL).

